# Lysenin-Induced Membrane Curvature on Erythrocytes and Nanoporous Mica Free-Standing Membranes

**DOI:** 10.64898/2026.11.05.764486

**Authors:** Elodie Lafargue, Natalia Martin-Gonzalez, Akiko Yamaji-Hasegawa, Toshihide Kobayashi, Adrian Carretero-Genevrier, Laura Picas, Ignacio Casuso

## Abstract

Beta porin toxins are virulence factors that oligomerize in host cell membranes, forming β-barrel pores that permeabilize target cells, leading to osmotic imbalance, cell lysis, and immune evasion. Despite extensive research on beta-pore-forming toxins (β-PFTs), significant gaps remain in understanding their role as actors of large-scale supramolecular membrane remodeling. While β - PFTs are traditionally viewed as passive cylindrical conduits, their biological efficacy is fundamentally tied to an evolved ability to structurally alter the host cell membrane. In this study, we combine erythrocytes, model membranes and a novel engineered nanopores on mica substrates. We overcome the limitation of traditional mica substrates and demonstrate that lysenin pores do not merely permeabilize the membrane; they curve it into rigid, dome-like supramolecular structures. Moreover, to gain biophysical insight beyond the standard membrane models which do not directly consider the membrane insertions, we propose a hybrid model combining the biophysical models and the architectural models of pendentive domes, where the intrinsic wedge-shaped geometry of the monomers is considered. Experimental mechanical assessment of the lysenin’s domes shows a non-linear mechanical of stiffness with respect the radius of curvature as predicted by our hybrid model. Collectively, our work suggests that the β-PFTs induce membrane curvature by the assembly of wedges. Beyond their pore-forming function, this membrane-bending activity may represent an additional mechanism by which β-PFTs interfere with host-cell membrane repair.

## INTRODUCTION

Membrane curvature drives essential physiological cellular processes including vesicle budding, endocytosis, cell division, and the functional spatial organization of membrane domains, among many others ^1–3^. Recent evidence suggests that membrane-disrupting pore-forming toxins (PFTs) respond to membrane geometry ^4^, and could also modulate it ^5–7^.

PFTs are primary factors for diverse pathogens and innate immune defense, typically existing as soluble monomers in their initial state ^8^. Among these, the β-PFTs uniquely create a stable, non-lytic intermediate named prepore ^8^. Next, under specific conditions ^8–9^, the prepore undergoes a dramatic conformational transition to reach a lytic state, the pore, where a transmembrane β-barrel permeabilizes the host’s cell membrane causing the cells ionic leakage and osmotic swelling ^10–12^.

How do β-PFTs remodel membrane shape beyond their pore-forming activity? The relationship between PFTs and membrane curvature has attracted increasing attention in recent years. Current evidence indicates that membrane curvature may play an active role in PFT’s action ^13^. Techniques such as Cryo-electron microscopy (cryo-EM), Giant unilamellar vesicle (GUV) fluctuation analysis, Nuclear Magnetic Resonance (NMR) spectroscopy, and All-atom molecular dynamics simulations point that PFTs’ action is not limited to pore induced membrane permeabilization but also involves several membrane modifications, such as membrane blebbing, fusion, and aggregation ^13^. The mechanisms underlying PFT-induced membrane curvature remain largely unexplored, leaving a significant gap in our understanding of the physiological and pathological implications of membrane remodeling by these toxins.

To observe PFT-induced membrane curvature, the scanning electron microscopy (SEM) offers static snapshots, while the Atomic Force Microscopy (AFM) and High-Speed AFM (HS-AFM) provides molecular-level imaging, mechanical assessment and dynamics under physiological conditions. Despite the AFM providing functional conditions where the process of protein-induced membrane curvature bending could take place under the observation of the AFM, these studies have rarely imaged the steps of action of the bending of the membrane by the proteins. The reason being that when imaging membranes by AFM, they are typically deposited onto solid supports, such as mica, which disfavors curvature generation, as the support-membrane interaction forces are stronger than the curvature-generating forces of the proteins. In fact, numerous AFM studies on PFTs behavior have only visualized the PFT action on a flat membrane geometries, hence, missing significant aspects of their behavior ^14–31^.

A novel mica substrate that overcomes the limitations of the mica-imposed flat geometry on the membranes was achieved. Fabricated mica nanopores wide of tens of nanometers and deep of tens of microns were created on mica. The freshly cleaved nanoporous mica is atomically clean of contamination and flat and the free-standing membranes are easily spread, contrary to the nanoporous supports used to date fabricated in silicon, alumina or gold ^32–37^.

Lysenin is a model for β-PFT studies ^38^, it has been extensively studied by AFM ^14–17^, but only on a flat-forced mica-supported geometry, so the curvature-related effects and mechanisms that the lysenin may exert on the membrane have been missed. Thanks to AFM it was revealed that lysenin increases its prepore to pore transition at low pH ^17^, or that the insertion in the membrane of the lysenin’s beta barrel alters the interactions and organization in the lipids in the membrane ^15^.

Lysenin specifically targets and binds to sphingomyelin (SM) in the membrane ^5^. The lysenin-induced membrane bending, if it was to take place, it would depends on the structural effects that the prepore and pore exert in the host cell membrane. Hence, the change of shape between prepore to the pore would have consequences in the membrane bending. Moreover, as the PFT tend to form dense oligomer assemblies ^39^, so the membrane-bending profile of each oligomer would be amplified (bending by crowding ^40^. In the existing models of protein’s membrane-bending, the continuum elastic Helfrich models ^41–43^, there is not a direct consideration of the protein’s shape, only an overall effective parameter accounting for the difference in curvature before and after the action of the protein, named the spontaneous curvature ^41^. To cope the limitation of the existing theory of continuum elastic Helfrich models, we introduce an architectural masonry-based model. In it, the β-PFT membrane-driven curvature is described using the universality of architectural building principles of bridges and domes ^44–46^. Our hybrid model considers specifically the disrupting bilayer integrity, where curvature emerges secondarily from pore’s shape in the membrane rupture, the supramolecular organization of the PFTs, and membrane-mediated protein-protein interactions. Such a theoretical framework combining architectural rules and membrane biophysics opens a new framework in the analysis of the bending of the membrane by proteins, specially indicated in the cases where the protein assembly forms dome-like structures.

Herein, we investigate the membrane-curvature generation by the pore-forming toxin lysenin. To maximize the relevance of our findings on lysenin action, we combined complementary experimental approaches to connect cellular relevance with mechanistic understanding: (i) electron microscopy (EM) imaging of living erythrocytes treated with lysenin, and (ii) topographic and mechanical atomic force microscopy (AFM) analyses of supported and freestanding membranes.

## RESULTS

### Lysenin induces temperature-dependent membrane curvature on erythrocytes

To check whether homo-oligomeric β-PFTs, as lysenin, can induce membrane curvature, in similar manners as it was identified for the hetero-oligomeric β-PFTs ɣ-hemolysin ^6^, we performed an *incellulo* assay using human erythrocytes. Moreover, to test the role of each of the lysenin conformers in membrane bending, we controlled the relative population of prepores and pores using temperature; cold temperature inhibits the transition to the pore state while maintaining membrane binding ^5^, hence we exposed the erythrocytes to lysenin for eight hours at 37 °C (dominant pore population) and 4 °C (dominant prepore population).

Prior to the toxin exposure, the electron microscopy images showed that the erythrocytes presented under control measurements using Bovine serum albumin showed an smooth cell surface (Fig.1a right). Following this first verification, the erythrocytes were exposed to lysenin at 100ng/ml (3 nM). First, we observed physiological conditions, at body temperature 37 °C, when the pores are assumed the dominant state, the erythrocyte surfaced showed a complete loss of smoothness over the totality of its cell surface. Over all the erythrocyte the SEM image showed the growth of protrusions of sizes ranging from tens of nanometers to a few hundreds of nanometers in diameter (Fig. 1.a, the circle of radius 200nm is a guide to the eye). At low temperature the effect of the lysenin on the erythrocyte presented a sharp contrast, 4 °C when the prepore state is assumed to be the dominant conformer and pore transition inhibited, the cell’s surface was smooth in most of its area and only at scattered locations the cell membrane was bent. Yet, the bending at those locations was very intense. Sharp conical protrusions with heights over 500nm and nanometric size apexes appeared scattered along the cell surface (Fig.1a second left, circle of radius 100nm; Fig.1a left, radius of apex protrusion ∼100nm).

**Fig. 1.**
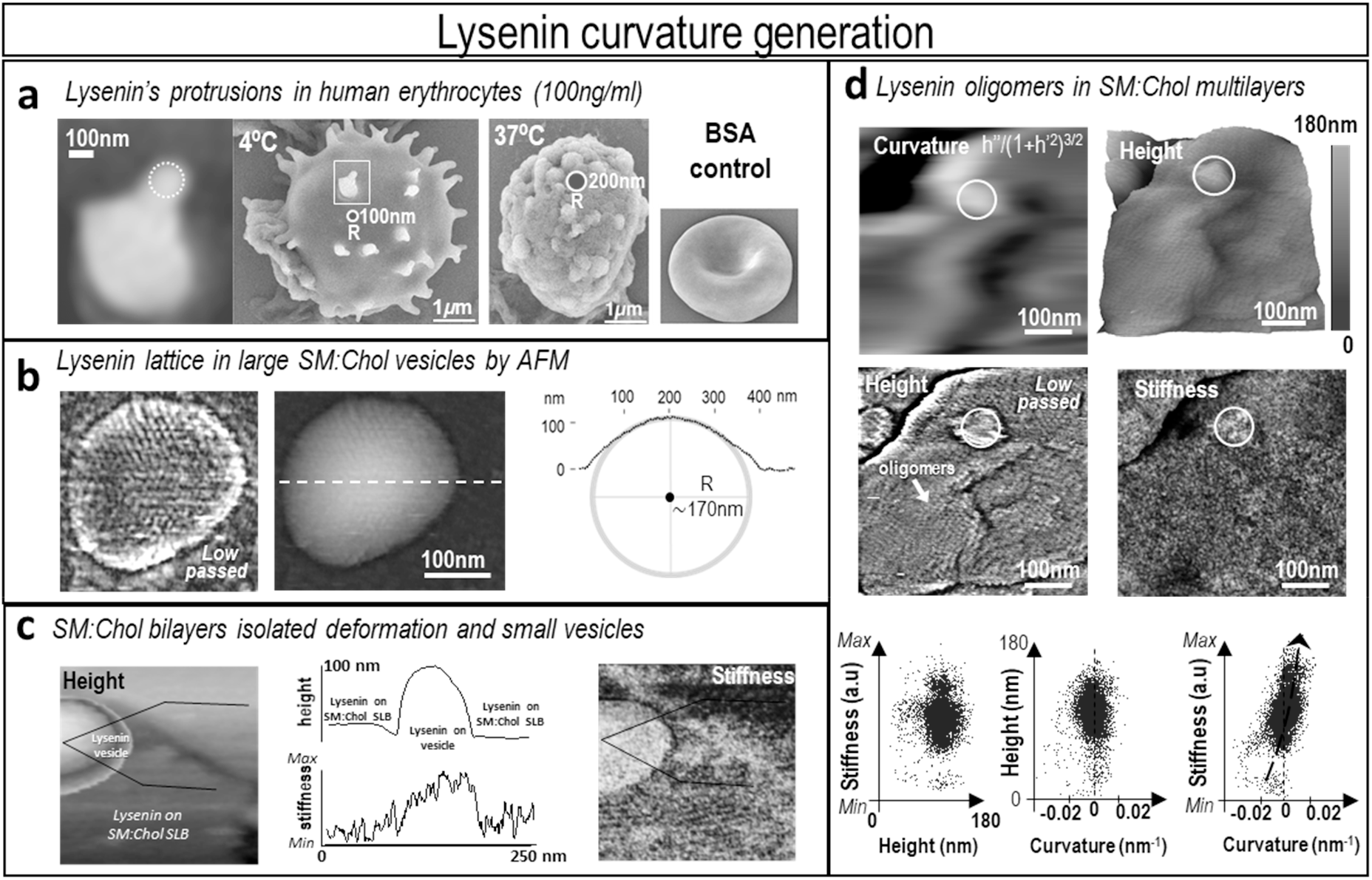
Lysenin-induced membrane curvature generation and nanomechanics. **a)** Electron microscopy of human erythrocytes exposed to 100 ng/ml lysenin at 4 °C and 37 °C. *Center:* Representative electron micrographs showing red blood cell morphology following lysenin treatment at 4 °C and 37 °C. *Right:* BSA-treated controls display no detectable surface deformation. *Left inset:* High-magnification view of the protrusion (indicated by the box in the adjacent panel) on red blood cells incubated with lysenin at 4 °C. Dashed circular guides highlight the protrusion curvature, corresponding to an approximate radius of 100 nm. **b)** AFM topography of a SM:Chol (1:1) vesicle coated with lysenin oligomers under pore-insertion conditions (pH 6.2). *Left:* Fourier low-pass filtered image revealing a well-or-dered hexagonal lattice of lysenin pores. *Right:* Profile analysis indicating a vesicle radius of curvature of 170 nm. **c)** Simultaneous topographical and mechanical map of a lysenin-coated vesicle surrounded by a lysenin-coated supported lipid bilayer (SLB) composed of SM:Chol (1:1). The mechanical map demonstrates significantly higher stiffness on the curved vesicle relative to the adjacent flat SLB. **d)** Topography and stiffness map of lysenin-coated SM:Chol (1:1) lipid multilayers featuring complex topography. Membrane curvature was calculated from the height profile using the first and second spatial derivatives along the X-axis. *Fourier low-pass filtered image:* Confirms complete coverage by lysenin oligomers. *Circled region:* Highlights a domain of high positive curvature (∩-shape) that exhibits correspondingly elevated stiffness. *Scatter plots:* Pixel-by-pixel correlations of stiffness-*versus*-height and height-*versus*-curvature show no systemic trend, whereas stiffness-*versus*-curvature reveals a positive correlation, where higher stiffness directly associates with increased positive curvature.

The results clearly demonstrate that the exposure to lysenin deforms the surface of the cells. The reason why it happens and it temperature-dependent manner is less clear. Possible causes include osmotic swelling ^47^ or the aggregation of lysenin in pore state, both causes are potentially be temperature-regulated, being affected by the phase transitions of the lipids in the membrane or the inhibition of the prepore to pore transition of lysenin. Other techniques are required to answer this open point.

### The hexagonal lattice of the lysenin pores accommodates the curvature in SM:Chol vesicles

To gain insight into how the lysenin organizes on and deforms the membrane, unilamellar vesicles of selected vesicular of diameters of hundreds of nanometers which mimic the size of the protrusions observed in the erythrocytes membrane were exposed to lysenin in solution at 10nM, in this condition the PFT populates and deforms without restrain the Sphingomyelin:Cholesterol (SM:Chol, 1:1) vesicles. To favor the formation of the pore state, the pH of the solution was set at 6.2 ^17^. After they were deformed, the vesicles were adsorbed in on freshly cleaved mica to be examined by AFM.

Topographical AFM images show that the lysenin had populated the vesicles with oligomers, most presumably in the pore stage, forming a dense hexagonal lattice of curved-shape, well-adapted to the round shape of the vesicles (Fig.1b left, Fourier low-passed). Inin this lattice the close-packed assembly was related to the membrane curvature, either adapting or causing it, suggesting that the pores insert in a shape of wedge in the membrane. Moreover, the AFM images showed that the radius of curvature of the close-packed assembly of lysenin pores was ∼170nm (Fig.1b centre). This value is in good agreement with that observed in the protrusions induced by the lysenin in erythrocytes, suggesting that sphingomyelin-bound lysenin may act as a scaffolding protein that generates and/or accommodates certain level of membrane curvature.

### Lysenin-coating, vesicles become stiffer than the supported lipid bilayers

The formation of a dense lattice of ‘wedges’ on top of a soft surface membrane can have significant impact on the mechanics of the membrane. The AFM uniquely enables the correlative assessment of mechanics and topography at the molecular level. To further investigate the behavior of lysenin-related membrane curvature, we performed mechanical characterization by off-resonance mapping^17^.

Under pore-insertion conditions (room temperature, pH 6.2 ^17^), the mechanics of the lattice of lysenin were characterized in two structural conditions: flat and curved. We located them in close proximity to enable simultaneous imaging to avoid uncertainties related to image to image acquisition instabilities (Fig.1c). The SM:Chol membrane, in its flat mica-supported lipid bilayer (SLB) configuration of height 6nm and in its vesicle configuration of height 90nm, was found equally densely populated by lysenin. Mechanically, the AFM found a surprising fact, contrary to the common situation in non PFT-coated lipid membranes, the stiffness was higher at the vesicle than at the SLB.

Commonly, the elevated rigidity of the mica substrate causes an effective stiffening of any supported sample, so the closer to the mica the stiffer the sample seems; a well-studied behavior known as the bottom-effect ^48^. Hence, commonly the SLBs appear stiffer than the vesicles. We verified that lysenin was was causing the unnormal behavior contrary to the bottom effect. To avoid uncertainties, in the same AFM image, we assessed, in a control assay, the stiffness of SLBs and vesicles of SM:Chol (1:1). We found (as expected) that the stiffness was lower in the vesicles than that in the SLBs (Fig. 2b), hence, confirming that the formation of a dense assembly of lysenin on the membrane modulates its curvature and the mechanics.

**Fig. 2.**
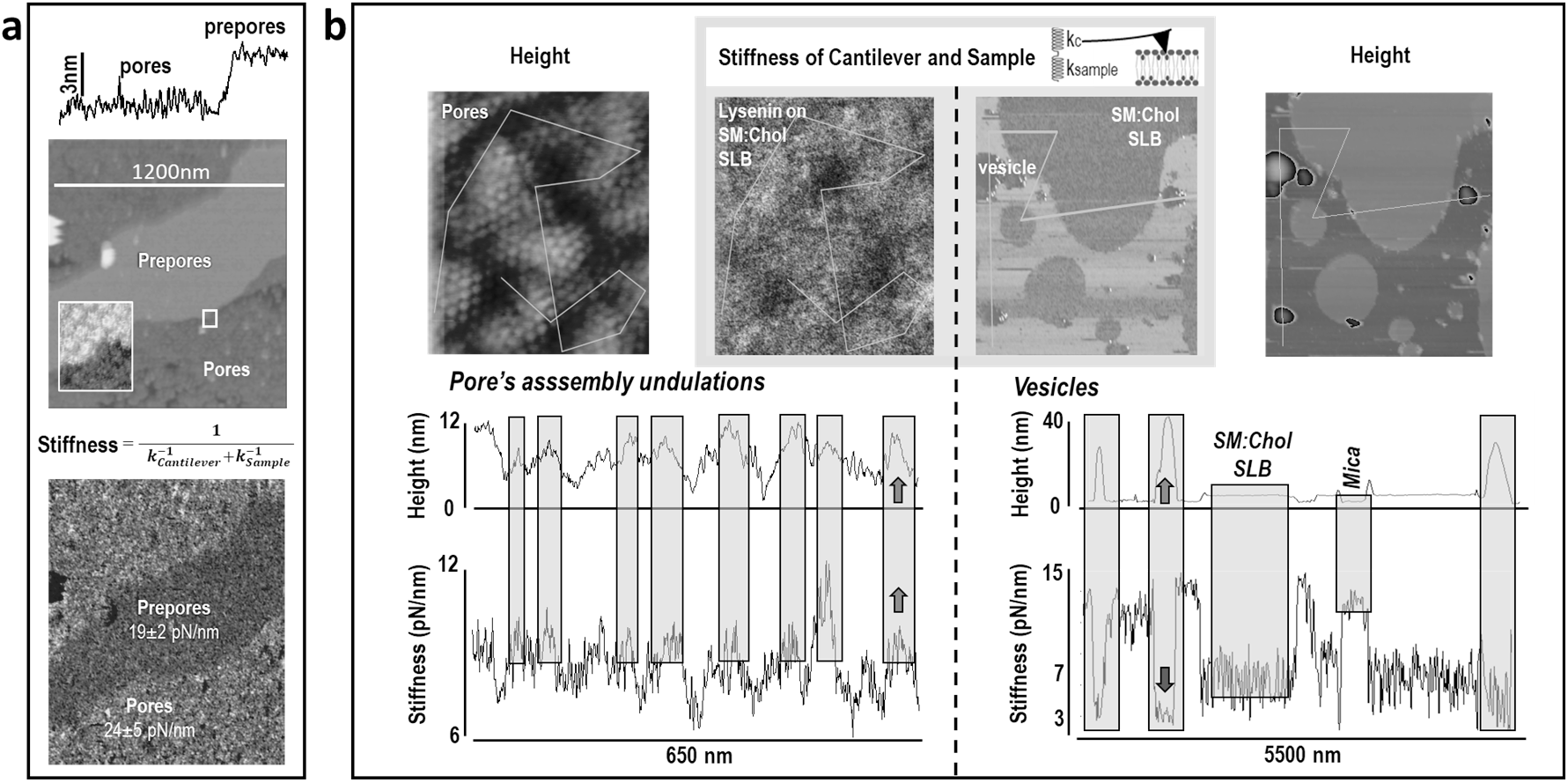
Extended incubation of lysenin on supported lipid bilayers reveals curvature-dependent mechanics. **a)** Mechanical map of a freshly cleaved mica substrate supporting a SM:Chol (1:1) lipid bilayer coated with lysenin following a 5 h exposure under mild insertion conditions (pH 7.2). *Height and stiffness images:* Topography reveals two distinct regions: an upper, flatter domain corresponding to lysenin assemblies in the prepore state, and a lower, rougher domain corresponding to lysenin in the pore state, with a step height difference of ∼3 nm. *Inset:* Close-up view of the oligomeric packing at the boundary between the prepore and pore phases. Based on the nominal cantilever spring constant (calculating overall stiffness as cantilever and sample springs in series), the average stiffness is higher in the pore region (24 ± 5 pN/nm) than in the prepore region (19 ± 2 pN/nm). **b)** *Left:* Mechanical map and height/stiffness cross-sectional profile of the undulated pore region. Topographical peaks on the undulations exhibit higher stiffness than the troughs, a mechanical trend that directly opposes the expected bottom-effect artifacts ^48^. *Right:* Control mechanical mapping on uncoated SM:Chol vesicles. The height/stiffness profile displays the opposite behavior, where the crests of the vesicles appear softer. The absolute stiffness decreases from the rigid mica substrate (stiffest) to the supported lipid bilayer (intermediate) to the bare vesicles (softest), consistent with standard bottom-effect behavior ^48^.

To further certify that our data is free of measurement error, rather than calculating the Young’s modulus, which relies heavily on contact area estimates (susceptible imprecisions in non-flat geometries ^48–49^, tip wear and contamination), we evaluated the system’s overall stiffness: force/displacement. The measured stiffness was the combined response of the sample and the cantilever, that can be modeled as springs in series (Fig.2, 1/[*k_Cantilever_*+*k_Sample_*]). Hence, when two samples were assessed in the same image, with the same probe, the measurement provides unambiguous comparison of their relative stiffness.

### Lysenin-coating, multilayered membrane’s stiffness becomes height- and curvature-dependent

Lysenin relates to membrane curvature, yet how it is not clear. To gain insight, we extended our previous assay that compared flat and round geometries to an assay richer in curvature environments. We generated rough surfaced multilayered membranes, whose folds create curved geometries ranging from positive to negative ^50–51^, and where, contrary to the SLBs, lysenin can deform the membrane free from the constraints of the mica substrate. Three rounds of adsorption of Giant Unilamelar Vesicles (GUVs) on a freshly cleaved mica surface, followed by exposure to lysenin at 10nM and pore-forming conditions (room temperature, pH 6.2), produced undulated membranes fully coated with lysenin oligomers (Fig. 1d).

Off-resonance mechanical mapping of the multilayers provided, besides the height and the stiffness, also the local curvature at each location of the multilayers. We calculated the local curvature of the membrane using the fast scan axis using the first and second derivatives of the height profile as follows (*κ*=ℎ′′/(1+ℎ′^2^)^3⁄2^), we selected a lateral step of the dimension of the lysenin oligomer 12.5nm ^14^, as it was justified biologically and showed relevant sensitivity. Such calculation provides assessment of the contribution of curvature to the mechanics of lysenin-coated membranes.

On the rough topography of the multilayers coated with lysenin oligomers, the stiffness, the height, and the curvature showed correlations in two types of spatial patterns (Fig. 1d). One, long range variations of the stiffness along hundreds of nanometers in the multilayers, could be due to the regions that changed in height without changing significantly their curvature. Two, in localized spots of tens of nanometers of highly curved membrane, which could be considered protrusions, the lattice of lysenin oligomers presented a particularly elevated stiffness (Fig. 1d, circle). Visual inspection shows the lysenin coating of the multilayers created specific and spatially discrete mechanics-bending relations. To gain further insight, we plotted the stiffness as a function of the height and curvature. The first finding was that there is not direct correlation between the stiffness and the height, all the range of stiffness was found at the same height (Fig.1d, middle right). The second finding was that the local stiffness showed a direct dependency on the curvature (Fig. 1d, bottom right): At negative curvatures (U-shape) the stiffness decreases, whereas at positive curvatures (∩-shape) the stiffness increases. The direct influence of the curvature on the stiffness is further supported by the lack of correlation between height and curvature (Fig. 1d bottom left). Overall, this data on the multilayers of SM:Chol showed that the lysenin related curvature modulates the membrane stiffness.

Unfortunately, the undulated multilayers do not allow to distinguish and certify whether lysenin oligomers are in the prepore or pore state using AFM topography. The decrease in height during the prepore-to-pore insertion is around 3 nm (i.e. slope 3nm/12.5nm = 0.24). This change is not detectable in the steeper slopes of the undulated membranes, which can reach 1 nm/nm.

### Lysenin pores induce the undulation and stiffening of supported flat membranes

Which of the conformers of lysenin is related to the curving of the membrane? Can the lattice of lysenin oligomers bend a flat membrane? To try to answer these points we developed a protocol on mica-supported SLBs based on the hypothesis that at sufficient slow-rate the lysenin could deform a mica supported SLB. We exposed the SM:Chol membrane to lysenin for 5 hours at pH 7.4, which is an intermediate strength of prepore-to-pore transition. Inspection by AFM showed that the lysenin oligomers segregated in space in two distinct conformer populations, each one presenting characteristic topography and also different stiffness (Fig. 2a).We identified two types of membrane surfaces (Fig.2a top): a smooth region containing flat prepores, with a RMS roughness of 2.4±0.5nm, and an undulated region containing inserted pores, with a RMS roughness of 4.9±1.2nm. The smooth prepore region was approximately3 nm higher than the roughed pore region. The height difference is consistent with that between prepores and pores ^17^, so we identified the smoother-higher region to the prepore conformer, and the lower-undulated region to assembled pores. The mechanical map revealed higher stiffness in the lysenin-pore region than in the prepore region (Fig. 2a, bottom), with spatially averaged stiffness were 24±5 pN/nm and 19±2 pN/nm, respectively.

After a 5-hour exposure of mica-supported SLBs to lysenin under conditions promoting a mild prepore-to-pore transition, we found that membrane-inserted lysenin bends the membrane. The finding that the pore layer is stiffer on average raises the question. Indeed, the pores sit approximately 3 nm closer to the mica surface than the prepores, and the bottom effect may therefore have contributed to their higher average stiffness ^48, 52–53^.

To rule out the potential contribution of the bottom-effect in the average stiffness we took a closer non-spatially averaged inspection. Detailed AFM images showed us that in the undulations of the pore assembly the relate between height and stiffness was contrary to that expected from the bottom effect ^48, 52–53^. The stiffness increased with the height (Fig. 2b, left); which proved that the stiffness in the membrane is not due to the substrate, but rather it is due to the specific mechanism of assembly of the lysenin pores that imparts structural rigidity to the undulating membrane. Consequently, regions of positive membrane curvature exhibit increased stiffness, whereas regions of negative curvature undergo softening.

The final proof that the lysenin is the determining factor that stiffens the membrane when it bends came from the mechanical assessment of vecicles and SLBs of SM:Chol in the absence of lysenin (Fig. 2b right). In these control experiments,performed in the same experimental configuration, the lower regions of the membrane are harder than the higher ones, as expected, if the lysenin is not present.

### Over nanoporous mica on freestanding membranes rigid lysenin domes are formed

Because the membrane repair mechanism of cells acts in seconds to minutes, our five hours assay may not be biologically relevant. To explore membrane deformation by lysenin at shorter timeframes, we used a novel type mica substrate, we fabricated nanoporous mica featuring pore diameters of 30 nm, our mica preserved the crystallinity and the depth of the pores enabled multiple rounds of mica exfoliation (Fig. 3a). So, its reusability, atomic flatness provided easy formation of supported lipid bilayers; a distinct advantage over other existing nanoporous substrate technologies ^34, 36, 54–55^. The nanopores covered 0.5% of the area of the mica surface.

**Fig. 3.**
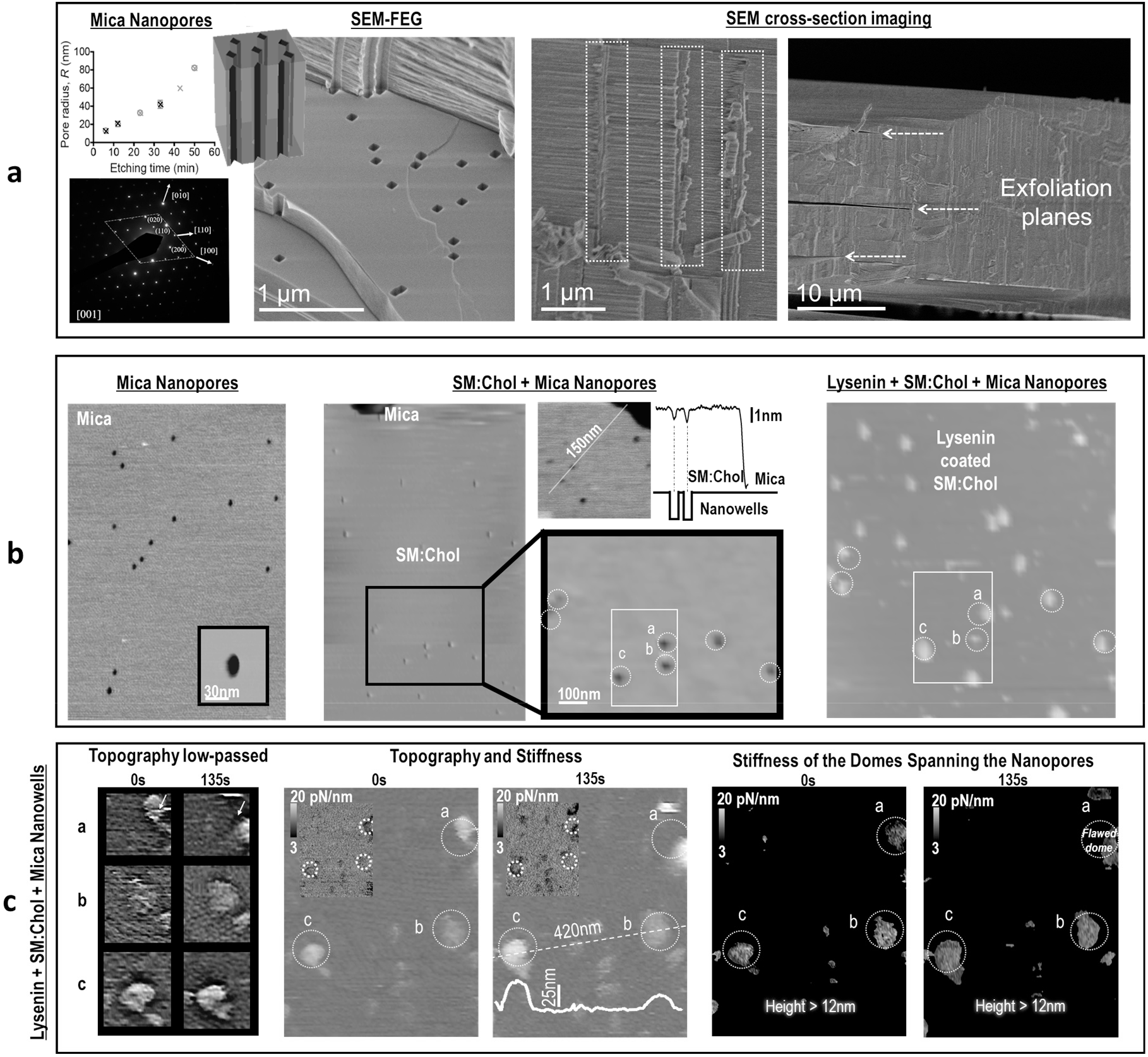
Lysenin-coated SM:Chol bilayers span nanoporous mica substrates to form structural domes. **a)** *Left:* 3D schematic model of nanoporous mica substrates. Etching time (5 to 55 min) controls the final radius of the nanopores. Electron diffraction confirms a monoclinic unit cell (a = 5.19 Å, b = 9.04 Å, c = 20.08 Å, beta = 95.5°) corresponding to the 2M1 muscovite structure ^76^, with rhombic pores (highlighted along the yellow dashed line) displaying high crystal order. *Center:* Transmission electron microscopy shows uniform nanopore size and spatial distribution, while cross-sectional views demonstrate straight, parallel nanochannels traversing several micrometers. *Right:* Demonstration that a single etched wafer can undergo multiple sequential cleavage cycles to generate multiple functional porous layers. **b)** *Left:* AFM topography of track-etched nanopores on a freshly cleaved mica surface. *Center:* Topography of an uncoated SM:Chol (1:1) bilayer spanning the nanopores alongside an exposed area of bare mica. The height profile reveals that the suspended bilayer sags slightly (∼1 nm) into the pore under the inward force applied during AFM imaging. *Right:* Upon coating the bilayer with lysenin under pore-insertion conditions (pH 6.2), rigid domes protrude outward above every nanopore. This dome formation across all nanopores is statistically significant, 6,500% more concentrated than expected under a random distribution. **c)** Temporal structural and mechanical stability of lysenin domes over nanopores. *Left:* Fourier low-pass filtered images confirming the presence of lysenin oligomer assemblies at t = 0 s and t = 135 s. *Center:* Topographical map of three domes spanning adjacent nanopores. *Inset:* Simultaneous stiffness map showing that intact lysenin domes exhibit local stiffness at their crests comparable to that of the lysenin-coated supported lipid bilayer on solid mica. While two domes remain structurally and mechanically stable throughout the 135 s observation window, the third dome is defective, showing temporal shape changes and markedly lower stiffness. *Right:* Masked stiffness map displaying only pixels corresponding to feature heights greater than 12 nm. The defective dome exhibits consistently lower stiffness compared to the two intact, stable domes.

We straightforwardly placed free-standing SM:Chol bilayer (1:1) on the nanoporous mica (see M&M). The chosen 30 nm pore diameter was more than three times larger than the dimension of an individual lysenin oligomer, enabling the assembly of multiple lysenin pores spanning the nanopore and, hence allowing lysenin to freely induce curvature in the bilayer.

The freestanding SM:Chol bilayers spanning the nanopores exhibited a minimal inward deflection of approximately 1 nm under forces applied AFM tip forces ranging up to few tens of piconewtons (Fig. 3b, center) ^56^. Hence, the free-standing bilayers did not undergo spontaneous deformation in the absence of lysenin.

When the freestanding SM:Chol bilayers were exposed to lysenin in pore-forming pH conditions and a oligomer saturating concentrations, we imaged by AFM mechanical mapping. Five minutes after exposure to lysenin, oligomers covered almost all the bilayer surface (97.6%) and appeared flat, reflecting the planar constraints imposed by the underlying solid mica support. However, non-flat lysenin-coated membrane geometries were observed on the remaining 2.4% (Fig. 3b).

Remarkably, the totality of nanopores covered by the SM:Chol bilayer exhibited a membrane bended by the lysenin overlying them. In terms of relative surface, only 2.4% of the membrane was bent, yet it located mostly at the locations of the nanopores with high statistical significance (Fig. 3b, dashed circles). Despite the nanopores occupying only 0.5% of the total surface area, roughly 33% of all bent membrane areas were positioned directly above mica-nanopores (Fig. 3b). This percentage is far above what would be expected from a random distribution: bending by the lysenin was approximately 6500 times [(33%/0.5% – 1) x 100] more likely to occur over a freestanding membrane region than over a supported region. Thus, lysenin-induced membrane bending preferentially occurs where the bilayer is free to deform, indicating that such bending is a natural response of the PFT when unconstrained deformation is possible.

The curvature induced by the assembly of oligomers showed a dome-like shape seen by the high-magnification AFM images (Fig. 3c, left), where the oligomers would present a wedge-like structural role. By using the AFM tip to assess the stiffness of the dome summits, it was found that the structural integrity of the domes constituted of lysenin wedges did was not dependent on the presence of an underlying mica substrate. The domes spanning the nanopores and those on top of the mica substrate presented similar stiffness (Fig. 3c), highlighting that the rigidity of these domes is exclusively determined by the structural arrangement of the oligomeric assembly, and not by the underlying mica support.

The mechanical maps revealed that the domes of lysenin have inherent structural variability. As certain domes showed higher mechanical fragility, that come together with a dynamic topographical reorganization over time (Fig. 3c, dome a, compare t = 0s to t = 135s). This mechanically-structurally instability could point to the presence of architectural defects in the domes, such as missing subunits or incomplete oligomeric packing, that would prevent a fully cohesive and rigid dome assembly.

All together, these findings suggest that the physical shape of the individual β-PFT oligomers, together with the fine details of their lateral interaction, directly determines the radius of curvature required to stabilize the dome structure governed by the wedge-like architecture of the lysenin pores.

### Hybrid architectural-biophysical modeling of lysenin’s domes: structural parallels to wedge masonry arches and domes

It was shown that pore-forming toxins like lysenin induce rigid dome-like protrusions in both erythrocyte and model lipid membranes. Structural and mechanical characterization by electron microscopy, AFM nanoindentation,and freestanding membrane assays demonstrated that lysenin oligomers assemble into lattices that create curved and rigid membrane architectures.Conventional continuum elastic models describe for protein-induced membrane deformation using phenomenological parameters, such as spontaneous curvature ^41, 57^. While useful in the long-wavelength continuum limit, such effective terms average out the discrete mechanical forces and localized contact stresses generated at the interfaces between individual protein subunits.

To uncover fundamental biophysical drivers governing the stabilization of these supramolecular assemblies, we integrated the lysenin pore shape and their associated inter-molecular force profiles into a cohesive mechanical model. A framework that provides a refined, quantitative interpretation of the experimental membrane deformations observed in vitro. Notably, the model revealed that individual lysenin monomers function as structural wedges operate under the same force-distribution principles as voussoirs and keystones in classical Roman masonry arches, as they maintain curvature and mechanical resistance against membrane tension.

Our model uses the load-path method and the voussoir force-balance principles used in architecture to study masonry Roman arches and vaults ^46, 58^. We rely on the fact that these mechanical rules governing compression-only stability and geometric alignment are scale-invariant ^44–46^. Our model links the shape and interactions of the toxin oligomers to dome’s shape and stability. We incorporated the event of the piercing and anchoring of the PFT into the lipid bilayer of the PFTs in the model, which is not accounted for in a direct manner in the continuum elastic models ^41–43^.

Like the domes of lysenin, the masonry domes and bridges are constructed by the repetition of identical wedge-shaped units forming dense assemblies, where the shape of the identical units determines at which curvature the construction withstands maximal loading. Moreover, as in masonry structures, the lipids between the PFT pores play a role analogous to that of mortar in bridges and arches: the mortar provides a flexible bonding between the wedges. Also, from architectural studies, we learn that the domes and arches become structurally unstable when their curvature deviates from the optimal stability radius *R_0_*. When R deviates from R0, adjacent wedges partially separate. This impairs load transfer across the contact interfaces and weakens inter-subunit binding, ultimately generating excessive curvature strain and structural collapse ^59–60^.

In our model, the lysenin oligomers and their closely associated lipids are the identical unit wedges that build the dome. The model assumes optimal/minimal energy bending if the pore-to-pore assembly is at maximum packing density. As a vertical force applied at the summit of the dome induces a displacement (*w*), the model determines the total dome stiffness (k_total_) through the addition of the mechanical response of the wedges and their inter-subunit connections (mortar). A calculation is achieved in three sequential steps: first, at each wedge to wedge contact, the distance between adjacent wedges (defined by the chord length, *L*) is related to the vertical displacement of the summit *w*; second, the interaction force or spring constant between adjacent wedges is calculated as it depends on the chord length *L*; and third, the contribution of all the wedges is integrated to calculate the total dome stiffness (*k_total)_*).

The integration of all the wedges was performed by the arch load-path approach. Widely used in architecture to obtain safe estimation of dome mechanics, in this approach the dome is represented as a set of meridional arches connected by hoop rings (Fig. 4a, right). The axial geometry of the force applied by the AFM probe enables the a load-path type of approach ^46, 61^.

**Fig. 4.**
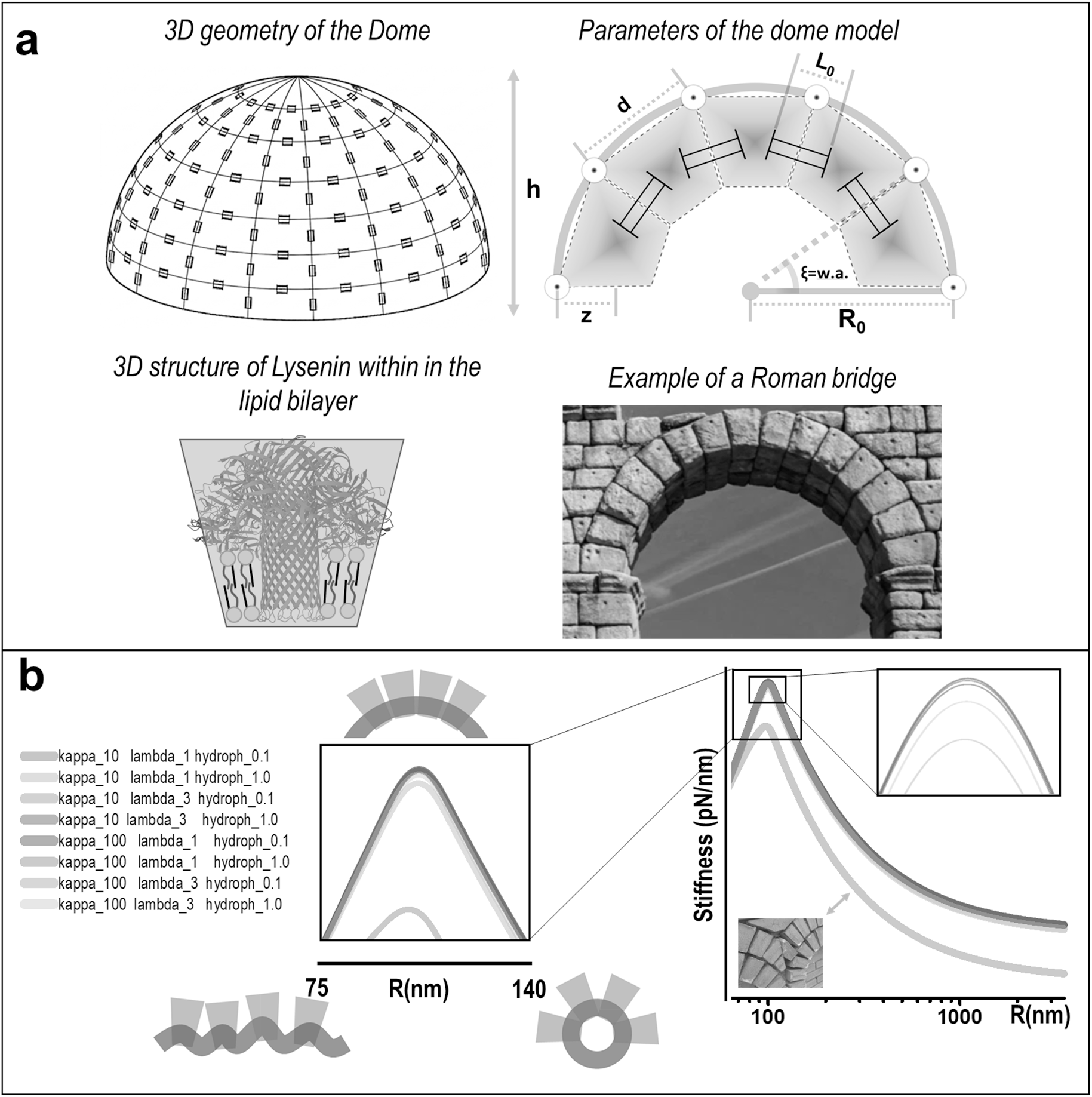
NanoDome wedge-and-mortar model of lysenin-induced membrane curvature. **a) Top left**. *Conceptual schematic and mechanical framework:* The collective deformation of the lysenin dome is modeled in response to applied vertical axial forces. The structural response arises from the joint mechanical action of adjacent protomer wedges. When adjacent wedges are not in physical contact, their interaction is described by the experimentally validated membrane-mediated protein–protein interaction profile, see 62. Upon mutual contact, the mechanical deformation of protein side chains and tightly bound lipid species is calculated using the Hertz contact model. Based on these individual wedge-towedge interaction profiles, the model calculates the mechanical force/deformation relationship for the entire dome assembly. Paths of axial load transfer are modeled as springs in series, while adjacent load paths are integrated as parallel springs 46. When vertical axial displacement takes place, the model calculates the changes in the distance between adjacent wedges. The change in distance gap for each wedge are derived geometrically under the assumption of equal angular deformation per wedge. The overall net axial stiffness of the dome as a function of its radius of curvature is obtained. **Top right**. The structural parameters of the model are depicted. **Bottom left**. 3D model of the lysenin pore, it is represented inside a wedge with associated lipids. **Bottom right**. A picture one of the arcs of the Roman aqueduct of Segovia, Spain, is shown to illustrate the parallelism between the architectural construction and the case of the lysenin. **b)** *Curvature-dependent stiffness profile and lipid sensitivity:* Deforming the dome away from its equilibrium radius of curvature (*R0*) causes a decrease in axial stiffness. For *R* > *R0*, vertical force flattens the assembly and reduces the stiffness; similarly, for *R* < *R0*, over-curving reduces stiffness too. Consequently, the dome exhibits a stiffness maximum at *R0*, which corresponds to the geometric state where the contact surface area between adjacent wedges is maximal. The variations in lipid bilayer properties cause the overall magnitude of the stiffness curve to shift up or down. We find that in the range of physiological values of lipid mechanics, there is minimal dependence of the dome’s structural rigidity on the lipid properties. Yet, there is a significant exception, brittle-like lipid properties (κ= 100kBT, decay length λ = 1 nm, hydrophobic mismatch = 1 nm; denoted by asterisk *) cause a pronounced reduction in axial stiffness, this decrease in mechanical integrity intensifies at larger radii of curvature (*R*). Mechanistically, these extreme conditions transform the lipid “mortar” from a flexible, long-range adhesive into a inelastic, short-range joint. The inset picture we that this behaviour has parallelisms in an arch falling out of place because the mortar was eroded causing the voussoirs to misalign.

To parametrize the dome’s architecture, five structural parameters were used: the dome radius *R*, the lattice pitch *d* defining the center-to-center spacing between lysenin pores, the wedge angle *w.a.* of the pore and its tightly associated lipid corona, the dome cap height *h*, and the hinge distance *z* measured from the outer dome surface to the central pivot point of inter-wedge load transmission (Fig. 4a).

The contact surface area between adjacent wedges reaches a maximum at the optimal dome radius *R_0_*, where neighboring faces align completely flush against one another. As the actual radius *R* deviates from *R_0_*, the rigid subunits are forced to pivot relative to one another about the hinge point, resulting in an angular mismatch and partial separation that systematically reduces the direct contact area while opening up non-contact gaps. To evaluate the net interaction force governing this dynamic interface, two contributions were added: (i) in the regions where the wedges do not contact, the long-range membrane-mediated protein–protein interactions operated, an exponential decay law was implemented, where the decay length *λ* is governed by the hydrophobic mismatch between the protein’s transmembrane domain and the lipid bilayer and the membrane’s bending rigidity *κ*; (ii) in the region of direct wedge-wedge contact, there is short-range mechanical resistance to compression, hence, the deformation was modeled by Hertzian elastic contact model ^62^.

The equations: The chord length between adjacent wedges is 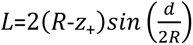. At the most stable geometry, it is (*L*_0,*eq*_) at 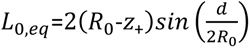, where 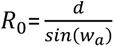. When the dome is deformed, the change of Chord Length (*ΔL*) with respect *R*_0_ is calculated by *ΔL*=*L*-*L*_0,*eq*_, and the variation of the pore-to-pore chord lengths with respect the axial displacement of the dome is given by 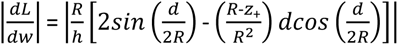. The force experienced in each unit is provided by the derivatives of the equations of the energy profile of the interaction between membrane inclusions with respect *L*. The total force per wedge-wedge interaction is given by an addition weighted to the contact and non-contact areas, which can be obtained from the cosine of the angle of the opening between the wedges. The force at compressed-contact area is 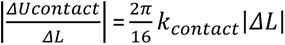, while at the Non-Contact Area (stretched regime) it is 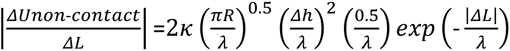. The stiffness (force/distance) at each wedge is calculated with respect the axial compression of the dome 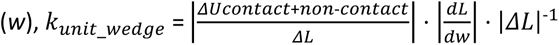. The dome is divided in arch load paths, where the total equivalent stiffness of each load path is a series of wedge units and it is calculated multiplying k_unit_wedge_ by a scaling factor provided by the pitch (*d*) divided by the arc length from the dome’s pole to its base, so 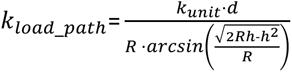. All the arch load paths are added, the total axial stiffness is calculated by multiplying *k_load_*___*_pat_*_ℎ_ by the number of load paths (*N)*, where *N* is approximated by the circumference of the cap’s base 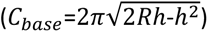 divided by the pore-to-pore pitch (*d*), resulting 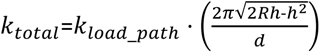

For the case of lysenin, we parameterized the architectural model using experimental structural data: a lattice pitch of *d* = 12.5 nm ^7, 11, 39, 63^; a dome cap height of *h* = 25 nm that matches the height we observed for domes spanning nanopores; an optimal curvature radius of R_0_ = 100 nm that is consistent with protrusions observed in erythrocytes (Fig. 1a), which gives a wedge angle *w.a*. = 0.12 rad; a hinge distance of *z* = 3 nm, that is one-third of the pore height which is the distance from the membrane’s hydrophobic core to the base of the cap; and an contact spring constant of *k_contact_* = 24.7 pN/nm (0.06 k_B_T / Å^2^), that we estimated from experimental HS-AFM data of interacting membrane proteins (ATP synthase) ^62^.

We tested the lysenin’s dome mechanical stability under axial compressive deformation (*w* > 0, corresponding to *R* > *R_0_*). When a vertical force is applied at its summit, we tested how membrane composition influences its stability. For that, we varied the membrane physical factors across their biologically relevant ranges. The bending modulus (*κ*) from 10 to 100 k_B_T ^64^, the hydrophobic thickness mismatch from 0.1 to 1.0 nm, and the interaction decay length (λ) from 1 to 3 nm ^62, 65–66^.

The dependence of the lysenin dome on the factors of the membrane was found weak in all the range of biophysical possible values. The model shows that lipid-dependent membrane properties shift the dome stiffness to higher or lower values. However, most of these lipid-related conditions only minimally alter overall stiffness across the analyzed physical range, regardless of the extent of deformation (at any radius R, Fig. 4b).

Interestingly, one specific combination of membrane conditions proves critical in degrading the mortar-like role of lipids that normally facilitates flexible bonding among pore-forming toxin wedges: when the membrane is highly rigid (*κ* = 100 *k_B_T*), the protein–protein interaction decay length is short (*λ* = 1 nm), and the hydrophobic thickness mismatch is large (1 nm). Under these specific bilayer conditions, the axial stiffness decreases significantly compared to other lipid environments. Mechanistically, these conditions turn the lipid “mortar” from a flexible, long-range adhesive into a brittle, short-range joint. As soon as axial compression forces the rigid wedges to pivot (*R* > *R_0_*), creating non-contact regions, the inter-subunit attraction collapses rather than providing a soft elastic restraint. This loss of attraction leads to the sharp drop in overall dome stiffness that becomes progressively more pronounced as compression drives the subunits further apart at higher *R* values (Fig. 4b).

Independent of lipid-dependent fluctuations, the dome always forms. The collective action of the wedges creates the curvature, and in all cases there is a defined curvature value where the coupling between adjacent pores provides the maximal structural integrity. This value corresponds to the geometry where wedge-to-wedge contact surface area is maximized. Consequently, the resistance to axial compression (dome stiffness) reaches its peak, hence an optimal radius of curvature *R_0_* (Fig. 4b).

The model finds a trend: the deformation away from the optimal radius of curvature consistently reduces the dome’s axial stiffness. When a compressive force applied at the summit increases the curvature radius (*R* > *R_0_*), adjacent wedges tilt apart and increase their inter-pore distance. Conversely, when a pulling force at the summit decreases the curvature radius (*R* < *R_0_*), the dome is forced into a tighter geometry that similarly reduces the contact area between adjacent protein wedges. In both regimes, the loss of direct subunit contact area systematically lowers overall dome stiffness (Fig. 4b).

The model showed that membrane bending mechanics are predominantly governed by the pore-forming toxins themselves. While lipid-mediated forces drive the protein crowding and can subtly modulate overall dome stiffness, the macro-level bending behavior of the membrane is ultimately dictated by the architectural geometry and structural rigidity of the protein wedges. This assembly behaves remarkably like classical masonry arches and domes, providing a mechanical explanation for how lysenin achieves such high structural stiffness when spanning nanopores on mica substrates.

Furthermore, the model identified that a planar configuration is not an intrinsically favored geometry in the dense assemblies of lysenin pores on lipid bilayers. In its pore state, the lysenin oligomer induces positive curvature in the membrane. Forcing the assembly into more planar geometries configuration, therefore loses its load-bearing efficiency, much like a collapsed Roman arch where the absence of proper curvature compromises load transmission and degrades overall structural integrity ^59–60^.

## DISCUSSION

While β-pore-forming toxins (β-PFTs) are frequently depicted as static, cylindrical conduits, their biological functionality is fundamentally tied to their capacity to actively deform and manipulate the host lipid bilayer. By relying on mica as a rigid, planar support, conventional AFM studies underestimate the membrane’s natural degrees of freedom, thereby masking the intrinsic capacity of lysenin to drive supramolecular membrane remodeling ^14–17^.

Investigations using planar supported lipid bilayers and AFM provided foundational insights into how PFTs assemble. However, these systems impose geometric constraints that limit membrane bending. To overcome this limitation, we combined a unique experimental platform track-etched nanoporous mica substrates with model membrane systems, and native erythrocytes to systematically map the curvature-dependent mechanical behavior of the β-PFT lysenin. By spanning geometrically defined bilayers and physiologically complex erythrocyte membranes, this approach enabled cross-validation of lysenin-induced membrane remodeling across complementary experimental systems.

Our multi-tiered approach confirmed that curvature generation is an intrinsic property of the aerolysin-family. By eliminating the physical constraint of the underlying flat substrate through the nanoporous mica architecture, we unequivocally demonstrated that lysenin oligomers do not merely reside within the bilayer. They actively remodel it into rigid, dome-like protrusions. We hypothesize that the asymmetric, mushroom-like shape of the lysenin oligomer may impose on the membrane a selective stiffening of the regions of positive curvature.

Our finding of lysenin-induced membrane curvature is in line with those on other PFTs. The ɣ-hemolysin was also found to induce pronounced membrane curvature ^6^. Yet, this is a bi-component bacterial β-PFT belonging to an evolutionarily distinct family, employing a different assembly mechanism (hetero-oligomeric pore formation). A convergence that suggests that the ability to remodel and bend host membranes may represent a widespread functional hallmark in the PFT families. Other PFT that has been identified with membrane curvature are the AB5 toxins (including the Shiga and cholera toxins) that employ it to facilitate cellular entry ^67^.

### A hybrid biophysical architectural model

We rationalized our experimental observations through universal architectural principles that transcend physical length scales ^44–45^, we established hybrid biophysical and architectural (wedge-and-mortar) model. By drawing an analogy between the hexagonal assembly of lysenin and an architectural pendentive dome, we demonstrated how the wedge-shaped geometry of the individual monomers produces a mechanically locked state. Treating β-PFT oligomers as structural wedges and the membrane-mediated protein–protein interactions as the mortar, allowed us to define an effective wedge geometry encompassing both the protein core and its tightly associated lipid corona. Crucially, our model revealed that the mechanical identity and overall stiffness of the curved membrane dome are predominantly dictated by the structural rigidity and shape of the β-PFT subunits, rather than the elastic properties of the lipid bilayer itself, which merely fine-tunes the dome’s mechanics.

These findings overturn the long-held assumption that PFTs passively adapt to lipid bilayer mechanics ^9^. Instead, the toxin forcibly re-shingles the membrane into a rigid architectural state whose axial stiffness is maximized at radius of curvature, *R_0_* ^44^.

At the structural level, the curvature generation may be coupled to the large prepore-to-pore conformational transitions characteristic of the aerolysin family, wherein the β-hairpin insertion loop unfolds to assemble the transmembrane β-barrel while the cap domain rotates away from the pore axis ^9^. We propose that during pore formation, the insertion of the β-hairpin acts as a mechanical lever, pulling on the outer leaflet to drive this structural evolution designed to lock and stabilize the pore.

From an evolutionary perspective, a lipid-independent and protein-dominated mechanism could confer a selective advantage on pathogens. It circumvents strict lipid-composition requirements, speeds up pore stabilization, and provides resilience against lipid-targeted host defenses, while ensuring precise geometry control and high mechanical robustness.

### Dual Cytotoxicity and Mechanical Subversion of Host Repair

Our finding of β-PFTs active membrane bending suggests that PFTs have a dual mechanism of cytotoxicity involving not only passive pore-mediated permeabilization, but also the active subversion of host cellular mechanics and membrane repair pathways ^68–70^. While eukaryotic cells typically counter PFT-induced permeabilization by shedding damaged membrane microdomains or internalizing them via endocytosis ^68–70^, the rigid physical constraints imposed by the lysenin oligomers present a potential mechanical blockade to these defenses. We hypothesize that lysenin actively impairs host cell repair machinery by imposing a rigid, outward dome-like topology. A negative curvature orientation that would directly oppose the inward, positive curvature required for classical endocytic invaginations, thereby repelling positive-curvature-sensing BAR domain proteins while selectively recruiting inverse-BAR (I-BAR) domain proteins ^71^. By physically locking the membrane in an outward deformation, the toxin would restrict the dynamic lipid plasticity required for membrane resealing, a mechanical arrest that would prevent an efficient bilayer repair, prolong intracellular ion leakage, and directly accelerate cell lysis ^72^. In addition, these reciprocal curvature-driven perturbations would likely disrupt downstream lipid signaling and membrane compartmentalization, particularly in highly active epithelial tissues enriched in sphingomyelin microdomains ^73^ or processes akin to clathrin-independent endocytic trafficking ^67^, and could amplify β-PFT virulence ^5, 16^.

In summary, our findings redefine β-PFTs from simple passive perforators into active agents of nanoscale mechanical remodeling. By utilizing nanostructured, freestanding nanoporous mica substrates and cross-validating across systems of increasing biological complexity, from synthetic model bilayers to native erythrocytes, we have uncovered an intrinsic, dome-like architectural curvature induced by lysenin assembly. We demonstrated that the structural rigidity of the protein oligomer overrides the intrinsic elasticity of the host lipid bilayer. The wedge-and-mortar mechanism indicated us that pathogens could have evolved to exploit membrane geometry as a physical weapon, effectively paralyzing host repair machinery and hijacking curvature-dependent signaling pathways ^74^.

Technically, beyond the immediate scope of pore-forming toxins, our development of freestanding lipid membranes on nanoporous mica provide a versatile biophysical platform for future investigations enabling easy examination of biomolecules-membrane interactions. The use of mica presents clear advantages over the rest of materials most commonly employed for the fabrication of free-standing membranes, such as silicon, alumina, silicon nitride or gold, as these are not reusable, lack atomic flatness, and get easily contaminated ^34, 36, 54–55^.

## METHODS

### Liposome formation and Supported lipid bilayer

Liposomes were prepared by the thin-film hydration method. Sphingomyelin (SM) and cholesterol (Chol) (Avanti Polar Lipids Inc., Alabaster, AL, USA) were mixed at a 1:1 molar ratio, dissolved in chloroform, and dried under a nitrogen stream to form a thin lipid film. Residual solvent was removed overnight under vacuum. The lipid film was rehydrated with 10 mM HEPES and 100 mM NaCl buffer (pH 7.4) preheated to 65 °C, followed by five cycles of 45 s heating at 65 °C and 45 s vortexing. The resulting multilamellar vesicle suspension was sonicated for 30 min in an ultrasonic bath. The liposome suspension was adjusted to 1 mg·mL⁻¹.

Supported lipid bilayers (SLBs) were formed on freshly cleaved mica. Prior to deposition, the liposome suspension was diluted 1:1 with imaging buffer (2 mM HEPES, 5 mM MgCl₂, 10 mM NaCl) adjusted to either pH 6.2 or pH 7.4. The diluted suspension was deposited onto the mica and incubated on a hot plate at 90 °C for 30 min to promote vesicle adsorption, rupture, and bilayer formation. The samples were then gently rinsed with the imaging buffer at the corresponding pH to remove excess vesicles and allowed to equilibrate in the HS-AFM liquid cell for at least 15 min before imaging. All HS-AFM experiments were performed in the corresponding imaging buffer.

### Lysenin protein

#### Purification and production

Lysenin with a *C-*terminal hexa-histidine tag plasmid were expressed in Lemo Competent *E. Coli*. Cells were grown in 2TY medium at 37 °C until reaching an optical density of 0.8 at 600 nm. Cells were collected by centrifugation at 4,000g and frozen until use. Cells were collected by centrifugation at 4,000g and frozen until use. Cell pellets were resuspended in 35 mM Tris HCl pH 7.6, 0.3 M NaCl, 2 mM TCEP, 5% glycerol and 10 mM imidazole (buffer A) before lysis by sonication. After, the soluble fraction was used for Ni-affinity purification on ӒKTA pure™ chromatography system with 5mL Cytiva HisTrap™ FF Columns. After, 3 ml of the pool Nickel purified from Lemo strain are dialysed O/N at 4°C in PBS. The dialysed pools were concentrated on 10 KDa concentrators. The final concentration is 0,5 mg/ml and stored in PBS pH 7.4 at -20 °C.

#### Protein incubation

To obtain a homogeneous assembly of Lysenin, 1 µM of Lysenin protein solution was directly deposited on a SM:Chol (1:1) SLB in a buffer of 2 mM HEPES pH 7.4, 5 mM MgCl2, 10 mM NaCl, incubated for 30 min at room temperature in a fume hood and rinsed with clean buffer 10 times. The sample was then mounted directly on the HS-AFM to perform the experiments.

### Nanoporous mica substrates

Nanoporous mica substrates were fabricated using a two-step procedure adapted from a previously described method ^75^. First, mica sheets were irradiated with high-energy heavy Xe ions (420 MeV) accelerated using a cyclotron (Cyclotron Research Center, Louvain-la-Neuve, Belgium, and GANIL, Caen, France). The use of heavy ions enables the generation of deep localized defects extending several tens of micrometers into the mica substrate. Second, following irradiation, the mica wafers were chemically etched by immersion in 20 wt.% hydrofluoric acid (HF, 11.4 M) for controlled durations. This mild etching process selectively enlarges the ion-induced defects, leading to the formation of randomly distributed rhombic nanopores. Pore dimensions were precisely controlled across etching conditions. Pore density reached an optimized value of 1·10⁹ cm⁻² (Fig. 3a). For instance, etching times of 25 min produced nanopores with short rhombic axis of approximately 30nm, as estimated from AFM topographic images (Fig. 3b). AFM characterization further confirmed the well-defined morphology and stable depth of the nanopores over multiple exfoliation cycles. Indeed, FE-SEM analysis of porous mica substrates confirmed a remarkable porous length over several tens of microns (Fig. 3a). Confirming a constant etching along the ion tracks. Structural analysis of nanoporous mica substrates was conducted using FE-SEM and TEM to verify crystallinity and measure pore dimensions (Fig. 3a). TEM and electron diffraction indicate that the mica surfaces are atomically flat, single-crystalline, and feature uniform pore diameters. Diffraction along the [001] zone axis reveals distinct crystallographic planes and the high crystallinity of rhombic pores (yellow dashed dots, Fig. 3a). The reconstructed unit cell is monoclinic, with parameters a = 5.19 Å, b = 9.04 Å, c = 20.08 Å, β = 95.5°, matching the expanded symmetry of the 2M1 muscovite structure ^76^.

### HS-AFM imaging and mechanical characterization

High-speed atomic force microscopy (HS-AFM) topographical and mechanical mapping was conducted at frame rates of 10–15 s/frame using a modified HS-AFM system (SS-NEX, RIBM, Tsukuba, Japan) equipped with BL-AC40TS probes (Olympus; nominal spring constant k = 0.1 N/m). To enable operation in Jumping Plus Mode ^77^, the standard control electronics were replaced with a custom controller developed by Nanotec Electrónica S.L. (Madrid, Spain).

In this mode, the probe executes vertical approach-retract cycles along the Z-axis at each pixel until a preset peak interaction force is reached. The tip then fully disengages from the surface before moving laterally to the adjacent pixel, minimizing lateral tip-sample shear forces. Topographical and mechanical data are acquired concomitantly by recording a complete force-distance curve at every pixel, yielding high-resolution local measurements of sample stiffness and adhesion.

Local stiffness was calculated during imaging by determining the slope of the repulsive contact region relative to the contact point. Stiffness values were normalized using reference force curves recorded directly on the rigid substrate to establish the maximum baseline stiffness across the topographical map, using the nominal cantilever spring constant.

#### AFM data analysis

Acquired image treatment was limited to the correction of XY drift and a first order X-line fit. Image treatment analysis and rendering was performed using the general distribution of WSxM ^78^.

#### Architectural model of the domes of lysenin

The modelling of the dome’s stiffness was carried out using Python code. The model integrates the mechanical properties of the proteins and the membrane-mediated protein-protein interactions ^62^ into a calculation method used in architecture for masonry bridges and domes, the arch load path calculation ^46, 61^. The code is available at https://github.com/DyNaMo-INSERM/Architectural-model-of-the-molecular-domes and also provided next:

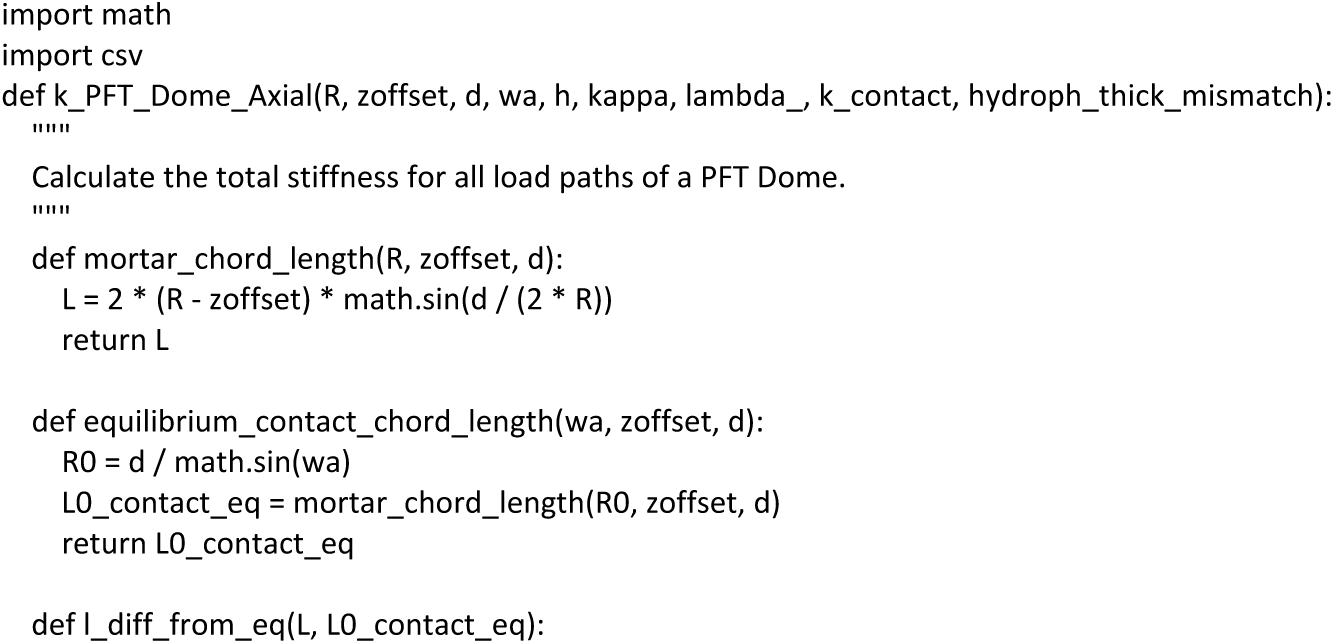
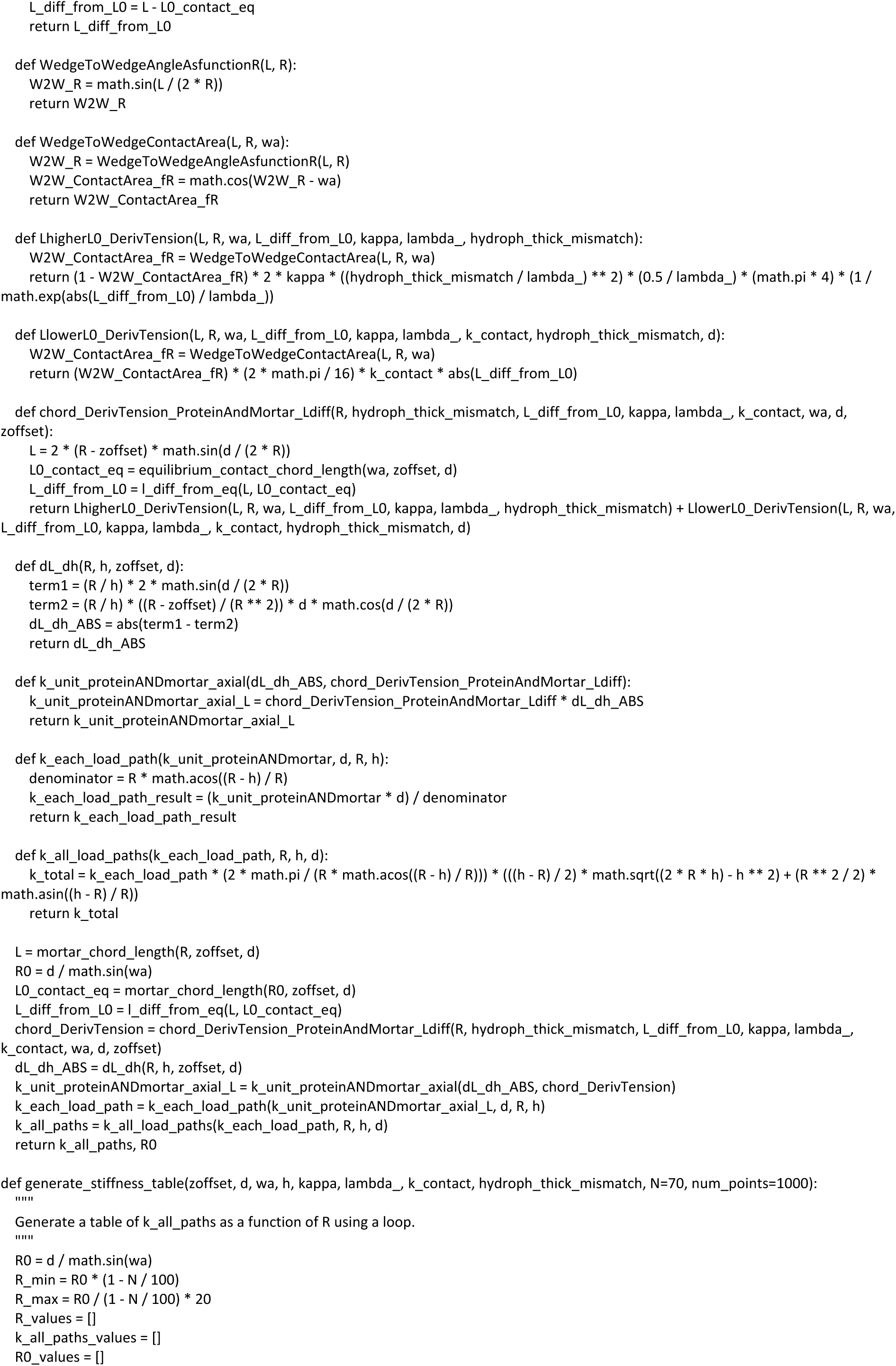
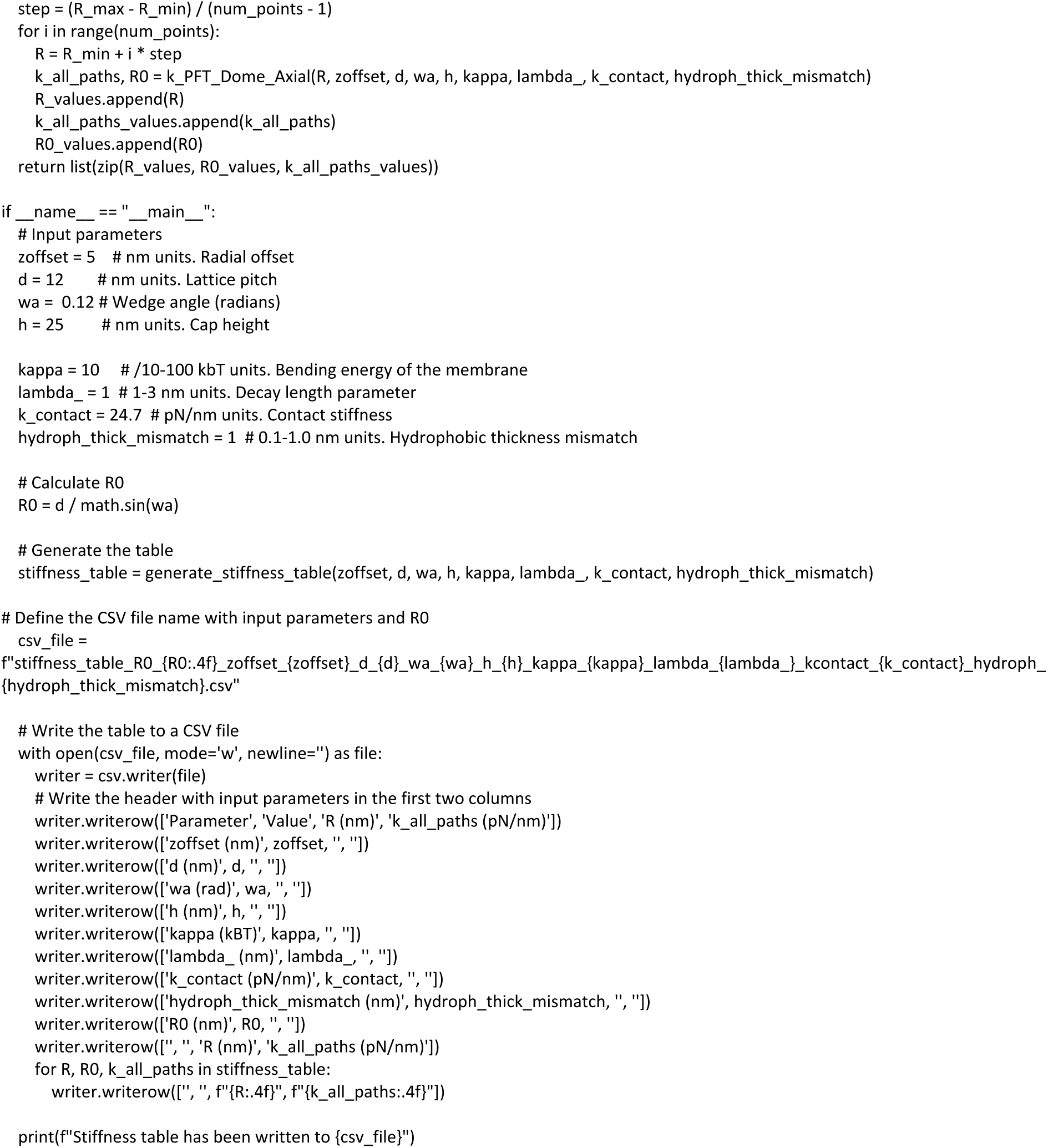

